# Learning differentially shapes prefrontal and hippocampal activity during classical conditioning

**DOI:** 10.1101/2020.12.02.408468

**Authors:** Jan L. Klee, Bryan C. Souza, Francesco P. Battaglia

## Abstract

The ability to use sensory cues to inform goal directed actions is a critical component of intelligent behavior. To study how sound cues are translated into anticipatory licking during classical appetitive conditioning, we employed high-density electrophysiological recordings from the hippocampal CA1 area and the prefrontal cortex (PFC). CA1 and PFC neurons undergo distinct learning dependent changes at the single cell level and maintain representations of cue identity during anticipatory behavior at the population level. In addition, reactivation of task-related neuronal assemblies during hippocampal awake Sharp-Wave Ripples (aSWR) changed within individual sessions in CA1 and over the course of multiple sessions in PFC. Despite both areas being highly engaged and synchronized during the task, we found no evidence for coordinated single cell or assembly activity during conditioning trials or aSWR.

## Introduction

The ability to react to sensory cues with appropriate behavior is crucial for survival. On the level of neuronal circuits, linking cues to actions likely requires the interplay between a large network of cortical and subcortical brain structures, including the medial prefrontal cortex (PFC) and the CA1 area of the hippocampus (Allen et al., 2017; Steinmetz et al., 2019). Both areas have been found to respond to sensory cues and reward in various behavioral paradigms (Aronov et al., 2017; Chen et al., 2013; Starkweather et al., 2018; Taxidis et al., 2020) and are involved in action planning and execution (Otis et al., 2017; Terada et al., 2017) PFC has been suggested to map contextual and sensory information to appropriate actions according to flexible rules (Euston, 2012). Accordingly, PFC has been found to control the development and expression of anticipatory licking during sensory guided reward-seeking behavior (Otis et al., 2017). PFC also maintains working memory representations of sensory cues over delay periods (Funahashi et al., 1993; Goldman-Rakic, 1995).

Similar to PFC, CA1 responds to sensory cues and displays sustained activity during delay periods to support memory formation (Hattori et al., 2015; McEchron et al., 1999; McEchron and Disterhoft, 1997).

Importantly, CA1 and PFC interact substantially during awake hippocampal Sharp-Wave Ripples (aSWRs) (Jadhav et al., 2016). aSWRs have been suggested to support planning of goal directed actions in the context of spatial navigation (Ólafsdóttir et al., 2018) and the disruption of aSWRs leads to impairments in anticipatory behavior (Nokia et al., 2012). In addition, sensory cue representations are reactivated in hippocampal and cortico-hippocampal circuits during aSWRs (Herzog et al., 2020; Rothschild et al., 2017). Yet, whether task related information during classical conditioning is also reactivated in the CA1-PFC circuit during aSWR and how this changes over the course of learning is currently unknown.

Here, we investigate how neural activity patterns in CA1 and PFC change throughout learning of sensory guided behavior, if information related to sensory cues is maintained while anticipatory actions are performed, and whether task-related information is reactivated in the CA1-PFC circuit during aSWRs. To this end, we employed high density silicon probe recordings from both areas in head-fixed mice during appetitive auditory trace-conditioning. Our findings reveal that CA1 and PFC exhibit distinct learning dependent changes in sensory cue evoked activity, trial type and sensory-cue related sustained activity as well as reactivation of task-related neural assemblies during aSWR.

## Results

### Head-fixed mice learn to anticipate reward during appetitive auditory trace conditioning

We trained head-fixed mice to associate one of two sounds (CS+ vs CS-) with a liquid reward delivered after a 1-second-long, silent trace period (Figure 1A) (Otis et al., 2017). Successful learning expressed as the emergence of anticipatory licking of the animals in response to CS+ sounds (Figure 1B & 1C). Across all animals, lick rates during the trace period were significantly higher during CS+ trials after 5 days of training (Figure 1C; Wilcoxon rank sum, p<0.01).

**Figure 1.**
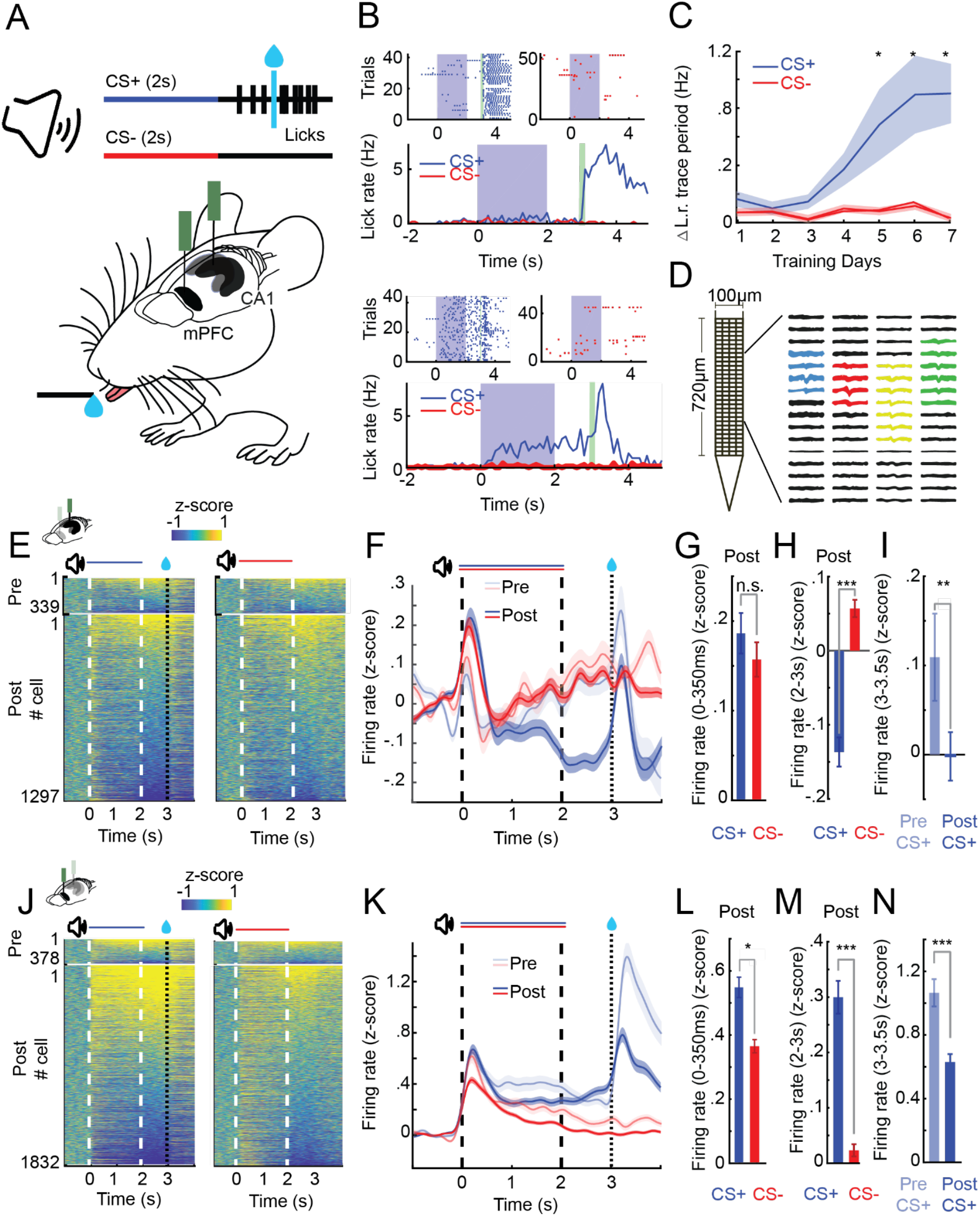
CA1 and PFC single cell activity shows distinct learning-dependent changes during appetitive auditory trace conditioning (AATC). **A)** Schematic of AATC task and electrophysiological recordings **B)** Example post-learning training sessions of one mouse during the AATC task (dots in raster-plots represent licks, solid lines indicate average responses from respective sessions). **C)** Average change in lick rate during the trace period trial during learning for all animals (n=17) (*indicates sessions with significantly higher group average licks during the trace period after CS+ sounds, Shade area represents standard error of the mean (SEM)). **D)** “Neuroseeker” silicon probe layout and combined spatial spike waveform patterns of 4 simultaneously recorded example neurons from CA1. **E)** Z-scored firing rates of all CA1 neurons recorded pre (top) and post (bottom) learning during CS+ and CS-trials ordered according to average trace period firing rates. **F)** Z-scored PSTHs of all recorded cells in CA1. **G)** Z-scored sound evoked change in firing rate (0-350ms post CS+/CS-onset) in CA1. **H)** Z-scored trace period change in firing rate (2-3s post CS+/CS-onset) in CA1. **I)** Z-scored reward period change in firing rate (0-.5s post reward presentation for CS+ trials pre and post learning) in CA1. **J)** Z-scored firing rates of all PFC neurons recorded pre (top) and post (bottom) learning during CS+ and CS-trials ordered according to average trace period firing rates. **K)** Z-scored PSTHs of all recorded cells in PFC. **L)** Z-scored sound evoked change in firing rate in PFC. **M)** Z-scored trace period change in firing rate in PFC. **N)** Z-scored reward period change in firing rate in PFC (*,**,*** represents Wilcoxon rank sum, p<0.05, p<0.01,p<0.001). (Error bars and shaded areas represent SEM).

### CA1 and PFC exhibit learning dependent changes in sound evoked and sustained activity

We next investigated how learning shapes neural dynamics in CA1 and PFC. To this end, we performed high-density silicon probe recordings from dorsal CA1 and PFC (1636 and 2217 cells total and 34 and 54 average per session in CA1 and PFC respectively; Supp. Figure 1 and Supp.Table 1), during the first two days of training and after the animals successfully acquired the conditioning task (hereafter referred to as pre and post learning) (Figure 1D, 1E and 1J). We observed that over the course of learning both areas developed pronounced differences in neural activity patterns between CS+ and CS-trials.

**Supplementary Figure 1.**
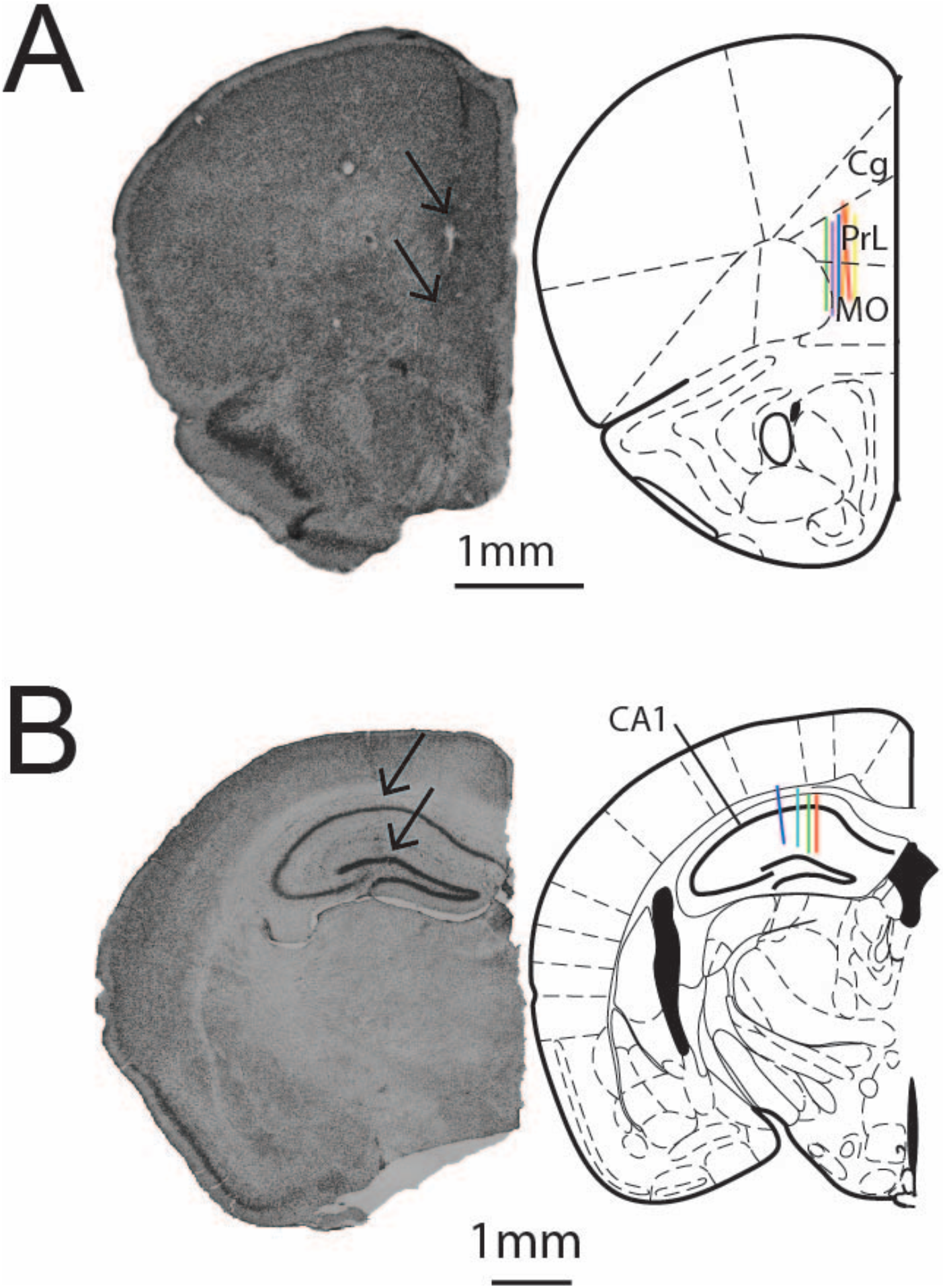
Positioning of silicon probes in CA1 and PFC. **A)** Histological image of silicon probe implantation track in PFC (left). Arrows show estimated extend of 128 channel “Neuroseeker” silicon probe. Schematic of verified recording position from 6 animals (right) (adapted from (Franklin and Paxinos, 2019). **B)** Histological image of silicon probe implantation track in CA1 (left). Arrows show estimated extend of 128 channel “Neuroseeker” silicon probe. Schematic of verified recording position from 4 animals (right)).

**Supplementary Table 1.** Number of recorded neurons per animal and session in CA1, PFC and simultaneous CA1-PFC recordings.

| *Animals* | **Pre Sessions** |  | **Post Sessions** |  |  |  |  |  | **All Sessions** |
| --- | --- | --- | --- | --- | --- | --- | --- | --- | --- |
| 1 | 15 | 14 | 43 | 25 | 64 | 56 | 36 |  | 253 |
| 2 | 40 |  | 85 | 53 | 19 |  |  |  | 197 |
| 3 | 20 | 24 | 54 | 37 |  |  |  |  | 135 |
| 4 | 28 |  | 52 | 39 | 41 | 42 | 42 |  | 244 |
| 5 | 31 | 29 | 34 | 39 |  |  |  |  | 133 |
| 6 | 36 |  | 34 | 33 |  |  |  |  | 103 |
| 7 | 31 |  | 31 | 22 |  |  |  |  | 84 |
| 8 |  |  | 83 | 42 |  |  |  |  | 125 |
| 9 |  |  | 31 | 41 | 38 |  |  |  | 110 |
| 10 | 79 |  | 83 | 80 | 77 | 7 |  |  | 326 |
| 11 |  |  | 62 | 66 | 62 | 65 | 50 | 69 | 374 |
| 12 | 57 |  | 77 | 94 | 48 |  |  |  | 276 |
| 13 | 68 |  | 34 | 22 | 84 | 41 | 47 |  | 296 |
| 14 | 42 |  | 44 |  |  |  |  |  | 86 |
|  | 69 |  | 22 |  |  |  |  |  | 91 |
| 15 | 17 |  | 34 | 40 | 36 | 42 | 16 |  | 185 |
|  | 76 |  | 49 | 22 | 64 | 16 | 66 |  | 293 |
| 16 |  |  | 29 | 40 | 20 | 12 |  |  | 101 |
|  |  |  | 14 | 61 | 35 | 52 |  |  | 162 |
| 17 | 12 |  | 44 | 20 | 12 | 27 |  |  | 115 |
|  | 29 |  | 16 | 32 | 43 | 44 |  |  | 164 |
| *Total* | 272 | 67 | 484 | 348 | 192 | 179 | 94 |  | 1636 |
|  | 378 |  | 471 | 460 | 451 | 225 | 163 | 69 | 2217 |

Short-latency evoked responses to both CS+ and CS-stimuli increased over the course of learning in CA1 (Kruskal-Walis test; F(3,3267)=23.64, p<0.001; Posthoc Wilcoxon Rank Sum Test for CS+, p<0.001 and CS-, p=0.008). In contrast, in PFC, CS+ responses remained high during learning but responses to CS-stimuli decreased (Kruskal-Walis test; F(3,4413)=21.49, p<0.001; Post-hoc Wilcoxon Rank Sum Test for CS+, p=0.2 and CS-, p<0.001).

During the trace period following CS+ sounds, cells in PFC exhibited, on average, a strong sustained increase in firing rates (Wilcoxon rank sum, p<0.001) (Figure 1M) while average single cell responses in CA1 became significantly suppressed (Wilcoxon rank sum, p<0.001) (Figure 1H).

Reward evoked activity significantly decreased in both CA1 (Wilcoxon rank sum, p<0.001) and PFC (Wilcoxon rank sum, p<0.001) (Figure 1I and 1N) from pre- to post-learning sessions.

### A subset of single cells in CA1 and PFC show lick related activity

Over the course of learning, mice started to respond to CS+ sounds with anticipatory licking. Therefore, we investigated whether CA1 and PFC also exhibited time-locked activity at the onset of anticipatory licking during CS+ trials. In line with previous reports showing the involvement of PFC in licking behavior during appetitive trace conditioning (Otis et al., 2017), we found that single cells in PFC showed increases in activity compared to pre-trial baseline at the time of the first anticipatory lick (PFC Lick Up, n = 77, 4% of all cells; Criterion: mean activity during -250ms - +250ms around 1st lick, 1 standard deviation above pre-trial baseline) (Figure 2D). We also found that a small population of CA1 cells responded during licking (CA1 Lick-up, n = 36, 2%) (Figure 2A).

**Figure 2.**
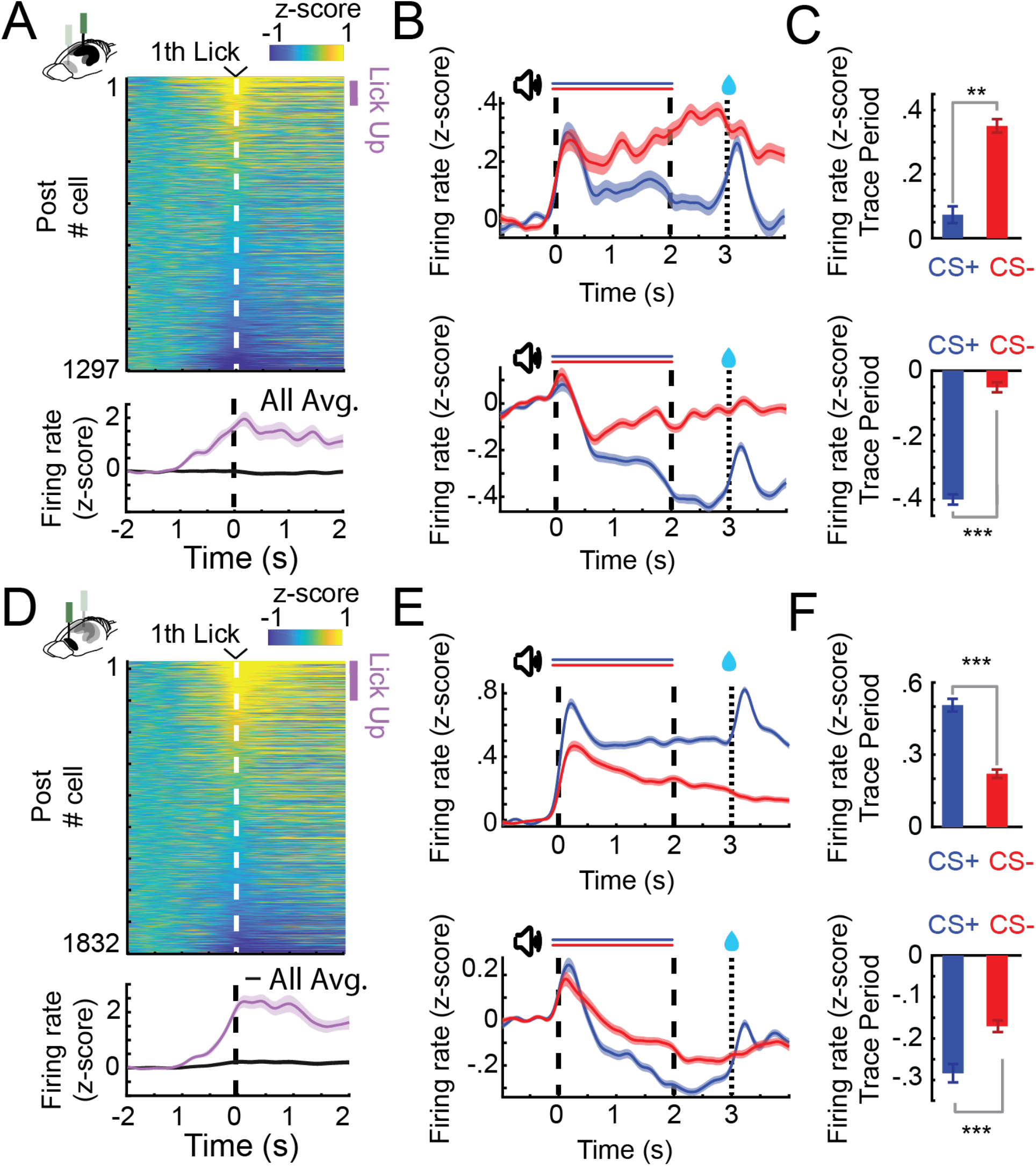
CA1 and PFC single cells exhibit lick evoked responses and distinct patterns of sustained activity. **A**) Z-scored firing rates of all CA1 neurons (Top) aligned to the first lick of a lick bout (at least 3 licks/s) during CS+ trials (before reward delivery). Z-scored change in activity of all and for positively lick modulated cells (bottom). Purple bar indicates Lick-Up cells. **B)** Z-scored PSTHs of all Trace-Up (Top) and Trace-Down (Bottom) post-learning for CA1 (in CS+ or CS-trials: Trace-Up, n=444; Trace-Down, n=675). **C)** Z-scored change in firing rate during the trace period of the same Trace-Up neurons (Top) and Trace-Down neurons (bottom) for CA1 **D**) Lick cells in PFC (same as in A) **E)** Trace-Up (Top) and Trace-Down (Bottom) non-lick neurons post-learning for PFC (CS+ or CS-trials: Trace-Up, n=736; Trace-Down, n=734). **F)** Z-scored change in firing rate during the trace period of the same Trace-Up neurons (Top) and Trace-Down neurons (bottom) for PFC. (*,**,*** represents Wilcoxon rank sum, p<0.05, p<0.01,p<0.001; Error bars and shaded areas represent SEM).

### CA1 and PFC cells exhibit distinct patterns of trial type-specific sustained activity

However, lick related activity was not the only driver of single cell modulation during the interval between CS+ and reward delivery. We discovered a large subset of single cells that exhibited trial type specific sustained responses during the trace period in CA1 and PFC, similar to what had previously been described during aversive eyeblink trace-conditioning (Hattori et al., 2015, 2014; Takehara-Nishiuchi and McNaughton, 2008).

In CA1, a large fraction of non-lick cells was significantly suppressed by CS+ stimuli (post Trace-Down, n = 580 40%) (Figure 2B and 2C (bottom)) and the percentage of these Trace-Down cells as well as suppression levels increased from pre to post learning (Pre 19%; Suppression Pre vs Post, T-test p<0,01). We also observed a smaller fraction of non-lick cells with sustained increases to CS+ sounds (post Trace-Up, n = 247, 17%) (Figure 2B and 2C (bottom)). The percentages of these cells decreased from pre to post learning session (Pre, 29%, Sup. Fig. 3 A & B).

In PFC, most modulated cells showed sustained increases in activity in response to CS+ sounds (Trace-Up, n = 734, 38%) (Figure 2E and 2F (top)) and the percentage and activation levels of these cells remained similar from pre to post learning. A smaller fraction of cells showed sustained suppression (Trace-Down, n = 630 33%) (Figure 2E and 2F (bottom)). In contrast, in CA1 differences between CS+ and CS-responses that emerged over the course of learning were mostly caused by a reduction in CS-stimulus evoked activity. CS-responsive Trace-Up cells in PFC significantly decreased their activity from pre to post learning sessions (T-test, p<0,001) (Supp. Fig. 2 A & B).

**Supplementary Figure 2.**
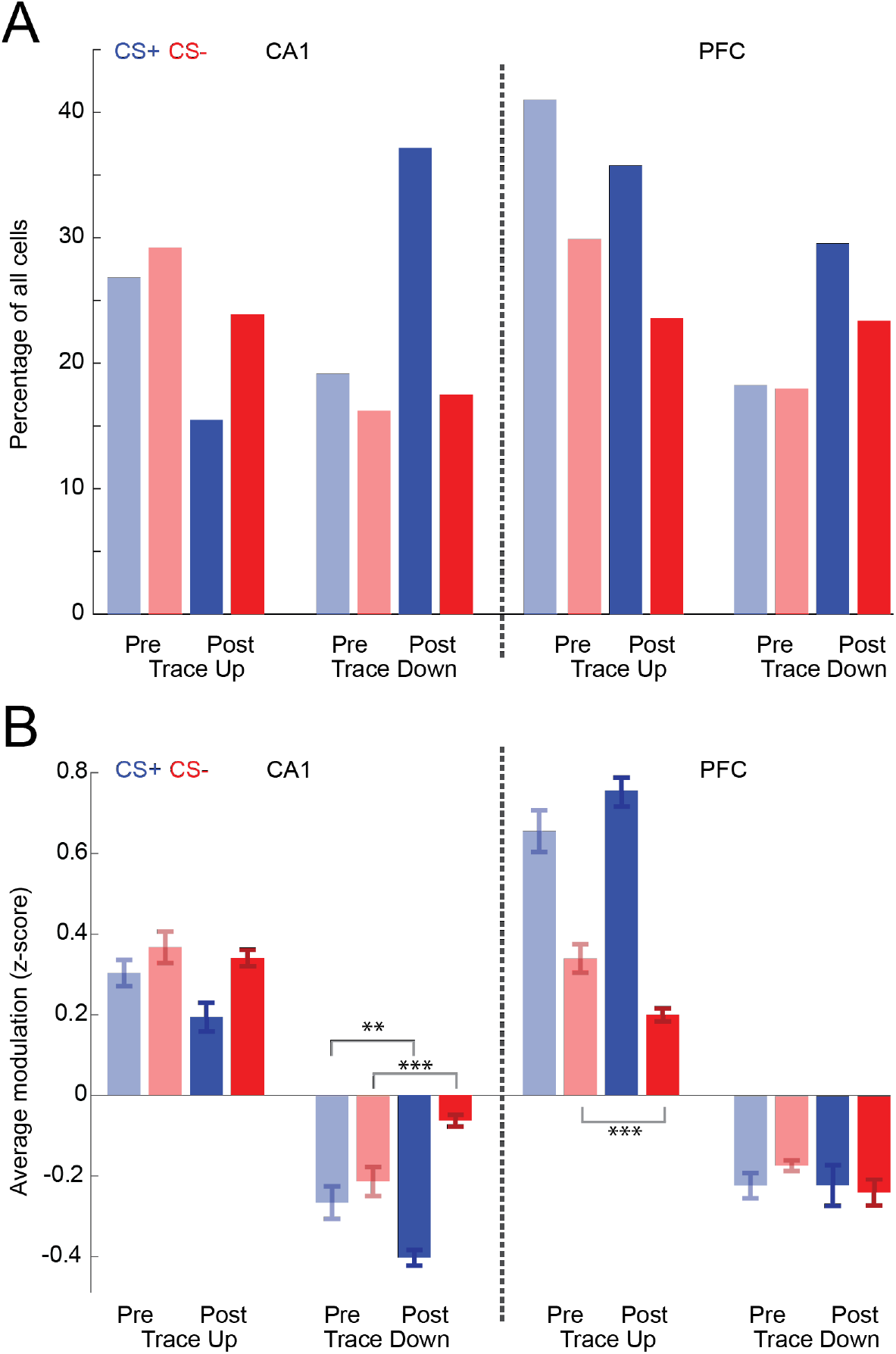
Distribution and activation of Trace-Up and Trace-Down cells in CA1 and PFC changes over the course of learning. **A**) Percentage of Trace-Up and Trace-Down cells in CA1 and PFC in pre- and post-learning sessions separately for CS+ and CS- trials. **B**) Average Z-scored modulation of all combined Trace-up and Trace-down cells in CA1 and PFC in pre- and post-learning sessions (Error bars represent SEM).

### CA1 and PFC population activity distinguishes between rewarded and unrewarded trials during the trace period

Because individual cells in CA1 and PFC exhibited sustained activity during the trace period, we hypothesized that these responses might be part of a broader CA1 and PFC population code to maintain a representation of trial identity between CS+ and reward delivery (i.e., in the trace period, in which there is no on-going stimulus). To test this, we first computed the binned population rate vectors for all simultaneously recorded non-lick cells in both areas during CS+ and CS-trials on a session-by-session basis. We next calculated the Euclidean distance between the population rate vector trajectories during the two stimuli (see Figure 3A and 3C for trajectory examples), and used this metric as a proxy for the similarity of population responses over time. We found that in post-learning sessions, the population rate vector distance increased after stimulus onset and persists to be significantly different from baseline during the trace period in CA1 (Wilcoxon sign rank, p<0.001) and PFC (Wilcoxon sign rank, p<0.001), indicating that both areas maintain trial type specific information at the population level. Non-lick cell population rate vector distance did not correlate with lick activity or movement of the animals in CA1 (n=36, Population Vector Distance vs. Licks: correlation coefficient =.14, p=.42; Population Vector Distance vs movement: correlation coefficient = .03, p= .85; Figure S3.2C) or PFC (n=38, Population Vector Distance vs. Licks: correlation coefficient =-.1, p=.52; Population Vector Distance vs movement: correlation coefficient = - .04, p= .74; Figure S3.2D). Population response differences persisted even after reward delivery and usually settled back at baseline levels after 20s (Supp. Figure 3.1).

**Figure 3.**
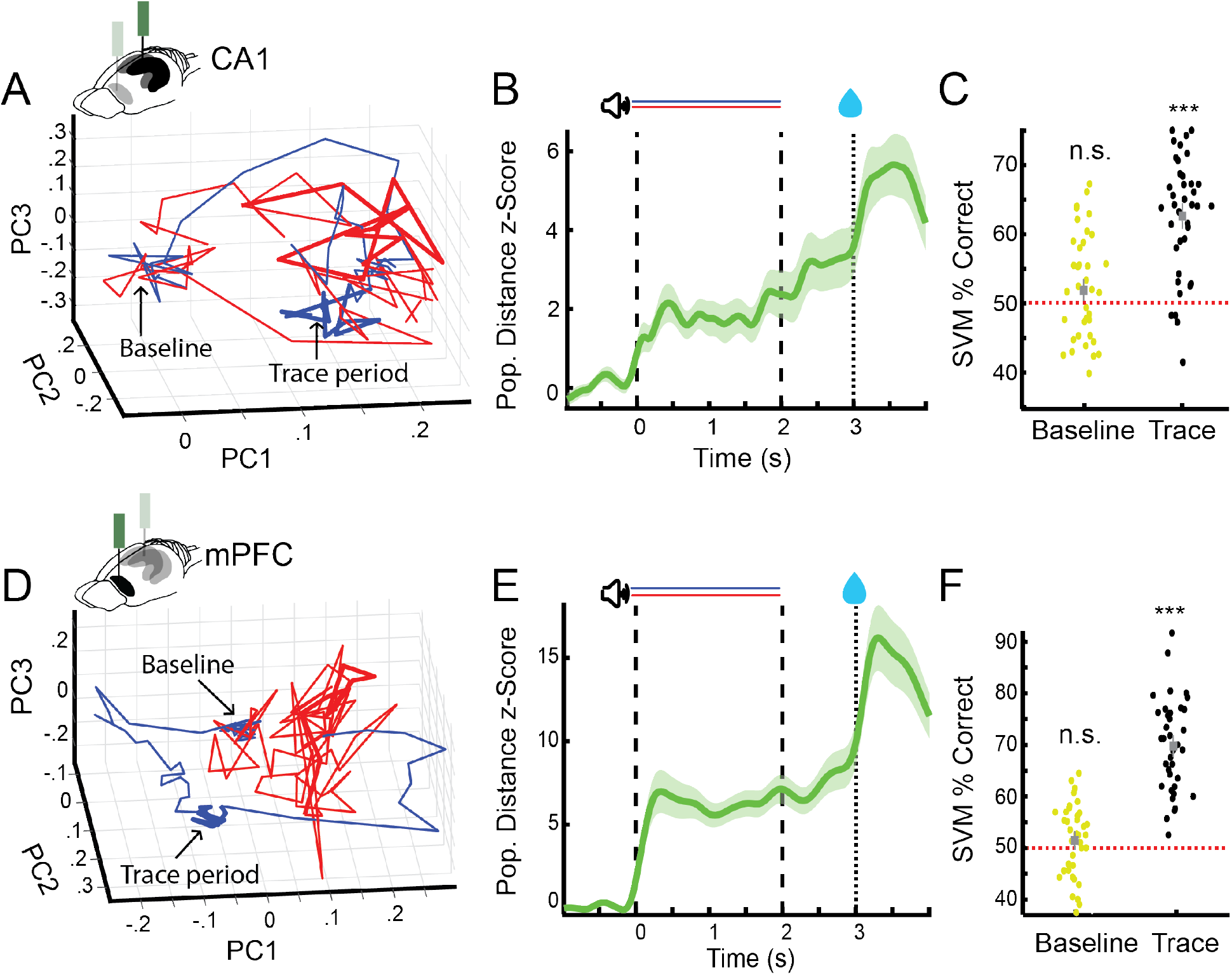
CA1 and PFC non-lick cell population activity encode trial identity during the trace period. **A)** Example of average non-lick cell population rate vector trajectories for one session in CA1 (CS+ (blue) and CS- (red)). Averages plotted along first 3 principal components (Baseline period: blob on the left, trace period thicker lines on the right). **B)** Average z-scored Euclidean-distance between CS+ and CS-non-lick cell population rate vector trajectories during AATC task for CA1 (n=36) (shaded areas represent SEM). **C)** Support vector machine classification of trial identity by average baseline (−1s-0) and trace period (2-3s) activity of non-lick cells in CA1 (n=36) (*** indicate Wilcoxon sign rank p<0.001). **D)** Example of average non-lick cell population rate vector trajectory for one session in PFC. **E)** Average z-scored Euclidean-distance between CS+ and CS-non-lick cell population rate vector trajectories for PFC (n=38) (shaded areas represent SEM). **F)** Support vector machine classification of trial identity by average baseline (−1s-0) and trace period (2-3s) activity of non-lick cells in PFC (n=38).

To verify that both areas maintain CS type specific information on a trial-by-trial basis, we next trained a support vector machine classifier on the firing rates of simultaneously recorded non-lick cells during the trace period from either CA1 or PFC. We were able to predict the preceding stimulus identity significantly above chance level in CA1 (mean performance, 62% correct, Wilcoxon sign rank p<0.001; Figure 3C) and PFC (mean performance, 76% correct, Wilcoxon sign rank p<0.001; Figure 3F).

**Supplementary Figure 3.1.**
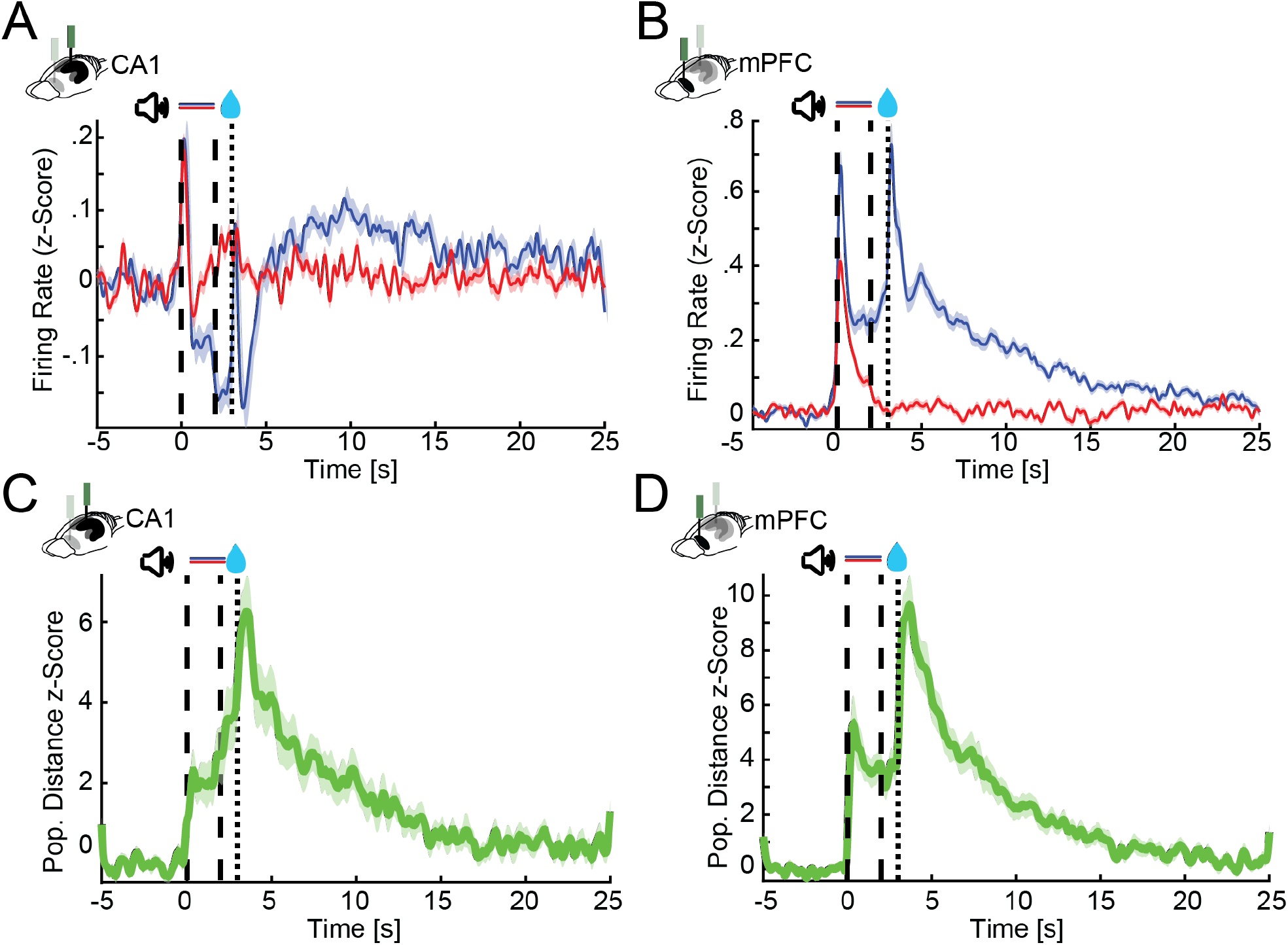
CA1 and PFC single cells and population responses slowly decay back to baseline after conditioning trials. **A & B**) Z-scored firing rates of all CA1 (A) and PFC (B) neurons recorded during post-learning sessions for 25s after trial onset. **C & D**) Average z-scored Euclidean-distance between CS+ and CS-non-lick cell population rate vector trajectories during AATC task for CA1 (n=36) and PFC (n=38) (shaded areas represent SEM).

**Supplementary Figure 3.2.**
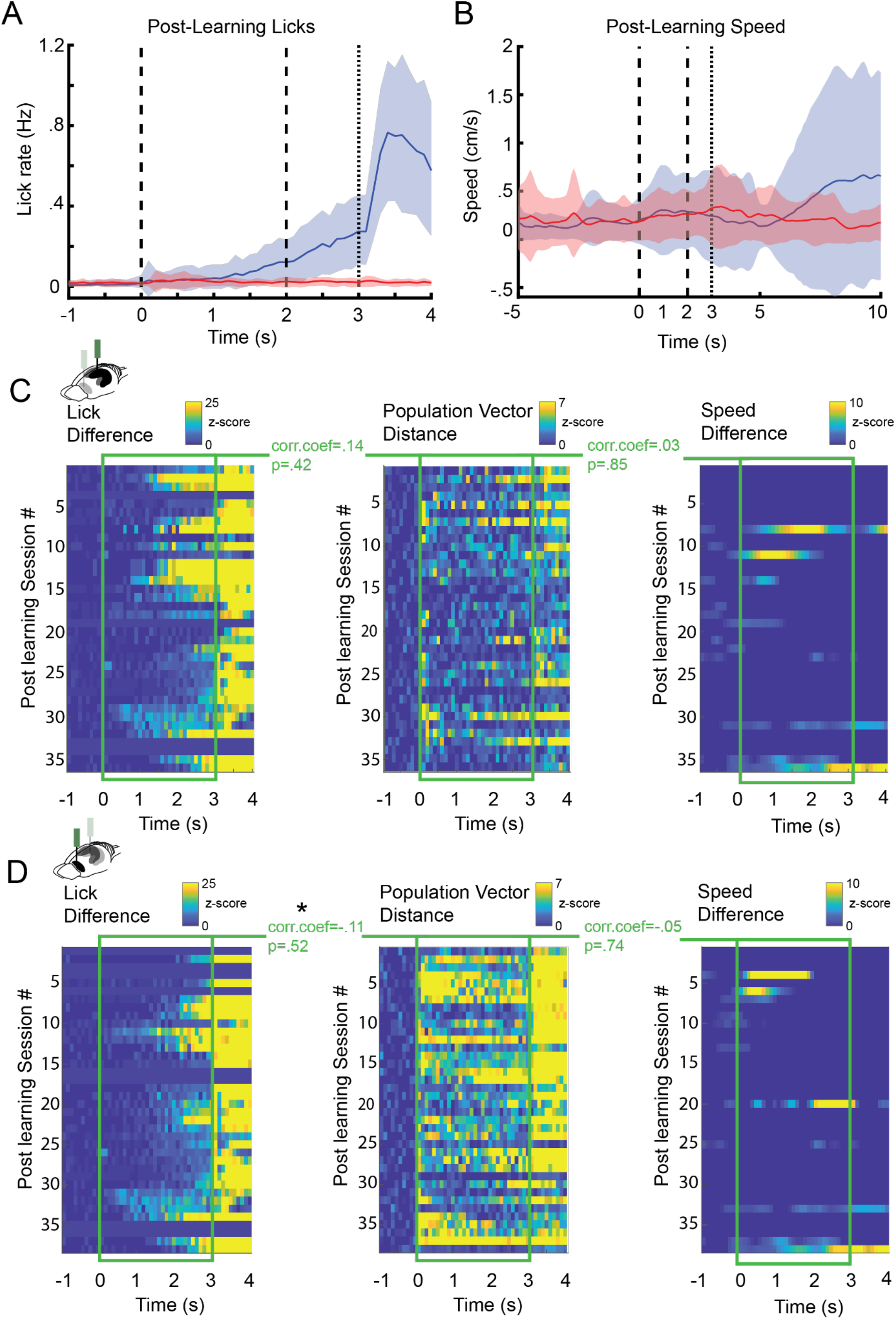
CA1 and PFC non-lick cell population activity does not correlate with lick or running behavior. **A)** Average Lick responses during post-learning trials for CS+ and CS-trials (52 sessions, 17 animals) (shaded areas represent SEM). **B)** Average running speed during post-learning CS+ and CS-trials (52 sessions, 17 animals). **C & D)** Correlation between non-lick population rate vector differences (CS+ vs CS-trials) for all post learning sessions in CA1 (C) and PFC (D) (middle) and differences in lick rate (CS+ vs CS-trials) (left) and differences in running Speed (CS+ vs CS-trials) (right) for all post learning sessions.

### CA1-PFC LFP coherence but not single cell interactions increase during trace conditioning

CA1 and PFC are known to interact during various spatial memory tasks in rodents (Benchenane et al., 2010; Jones and Wilson, 2005; Sigurdsson et al., 2010; Spellman et al., 2015) and aversive trace-conditioning affects local field potential synchronization within and across brain areas (Shearkhani and Takehara-Nishiuchi, 2013; Takehara-Nishiuchi et al., 2011). Given that CA1 and PFC are also highly engaged and encode information about trial identity during appetitive trace-conditioning, we wondered if we could find evidence for an interaction between CA1 and PFC on the level of single cells and local field potentials (LFP). We found that post-learning, high frequency CA1-PFC LFP coherence was increased during CS presentation (n=13, 60-140 Hz permutation test at each frequency <0.05) (Figure 4A). During CS+ trials, trace period coherence was furthermore significantly elevated across a broad frequency range (n=13, 12-140 Hz permutation test at each frequency <0.05).

**Figure 4.**
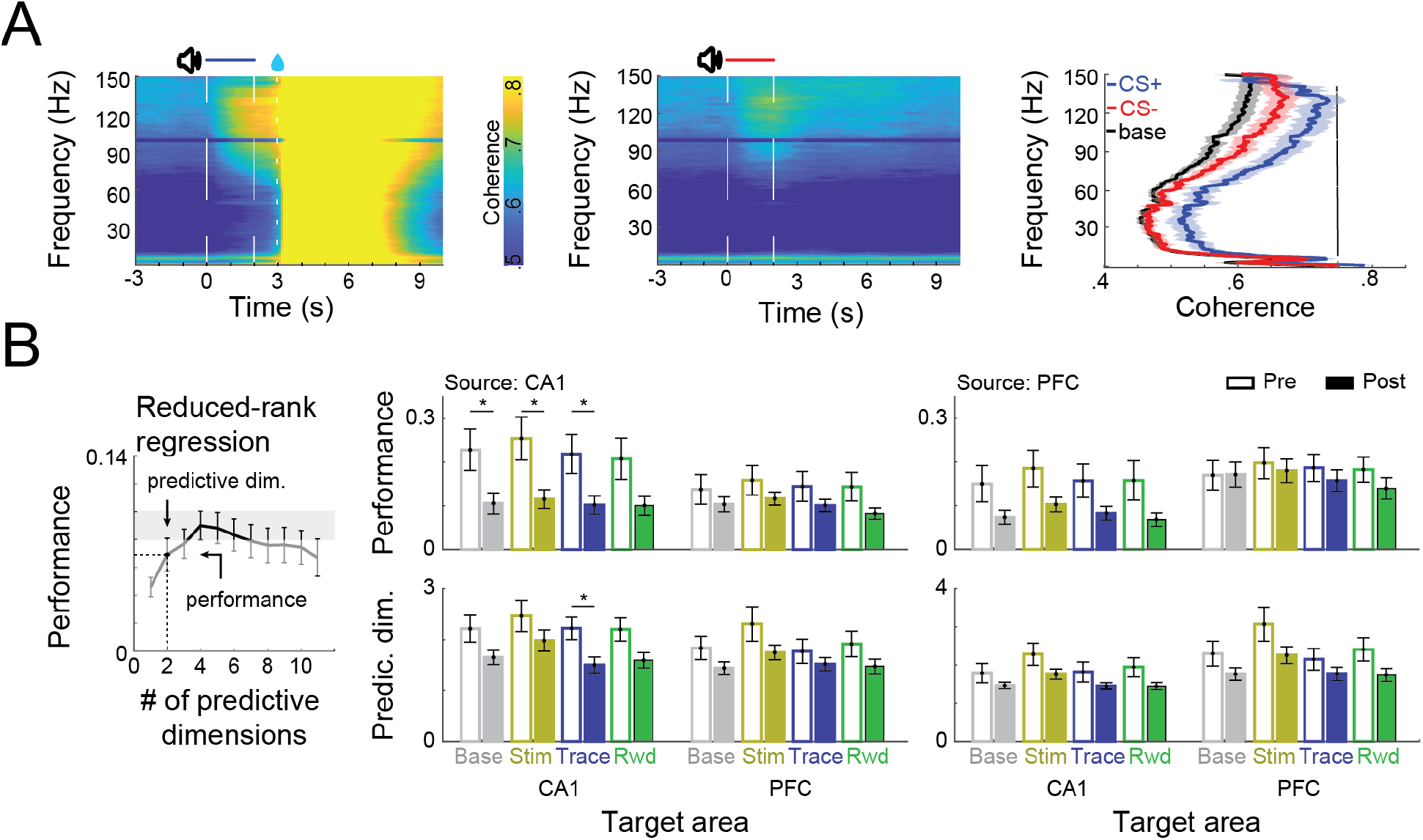
CA1-PFC interaction during trace-conditioning. **A)** CA1-PFC LFP coherence during CS+ trials (right) and CS-trials (left). Average coherence during baseline and during the trace-period (right). Black bar indicates significant difference between CS+ post and CS-post trials (permutation test at each frequency <0.05). **B)** (Left) Schematic representation of how performance and the number of predictive dimensions were calculated for each regression. (Right) Reduced-rank regression between CA1 and PFC spiking activity during conditioning trials in pre and post learning sessions. Solid and filled bars represent pre learning and post learning sessions respectively (error bars represent SEM).

Given the interactions between CA1 and PFC on the LFP level, we next checked for an interaction between both areas on the single cell level (Figure 4B). To this end we computed a reduced-rank regression (RRR) to assess how well the activity of a sampled population in one of the areas (source area) could be used to explain another (disjoint) sampled population in the same area or in another connected area (target area), through a simplified, low-dimensional linear model. We then used cross-validation to estimate the optimal dimensionality (rank) of each RRR and its performance (R^2^) (Semedo et al., 2019).

We observed that post learning CA1 ensemble activity at baseline could be used to predict other individual neurons firing rates in CA1 just as well as the firing rates of neurons in PFC. The PFC ensemble on the other hand was much better at predicting the firing rates of other PFC neurons compared to neurons in CA1 (Figure 4B), which indicates a directionality of information flow between both areas at baseline.

However, cross-area predictability of firing rates did not change significantly when we compared baseline levels to any of the different trial stages (Stim, Trace or Reward) (Figure 4B). Comparing the performance of the full-rank model also did not reveal any significant differences in coordination between CA1 and PFC across task periods (Ridge-regression with L1 regularization; Supp. Figure 4)

However, by focusing on CA1, we found that ensemble activity substantially decorrelated over the course of learning and individual cells firing rates were significantly less well predicted by the rest of the ensemble from pre to post learning sessions. This was not the case in PFC (Figure 4B).

**Supplementary Figure 4.**
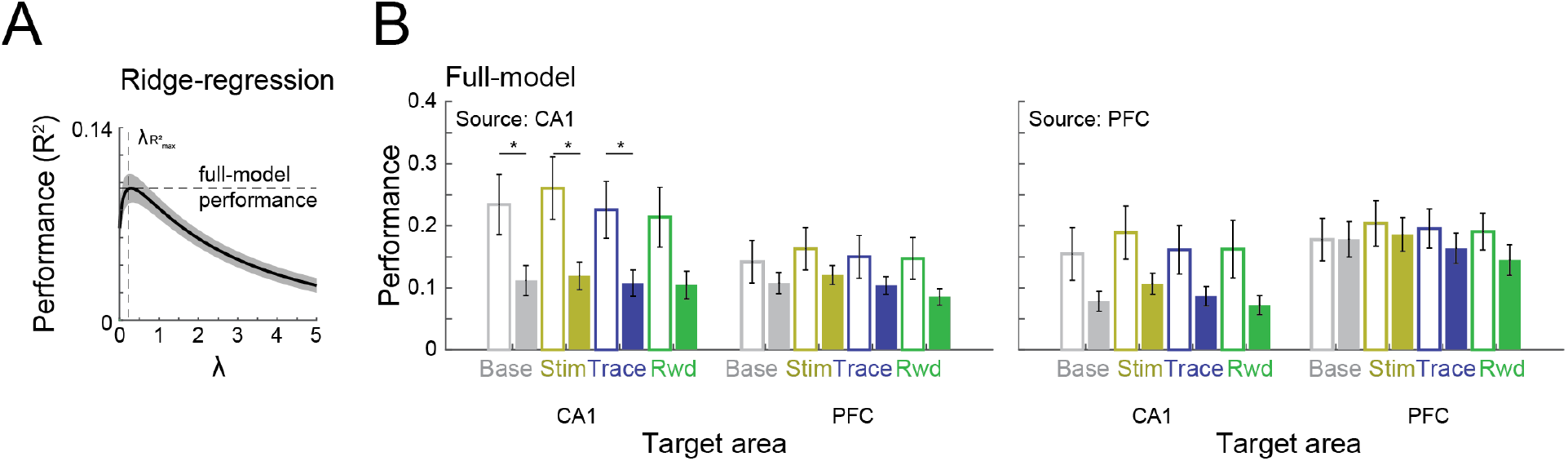
CA1-PFC single cell interaction does not change across different task periods. **A)** Schematic representation of Ridge-regression. A full-rank model was computed using 10-fold cross validation and L1 regularization. The model with the best performance over the regularization parameter λ was selected. **B)** Full-model ridge regression between CA1 and PFC spiking activity during conditioning trials in pre and post learning sessions. Solid and filled bars represent pre learning and post learning sessions respectively, error bars represent SEM, and * refers to p<0.05 in a Wilcoxon ranksum test.

### Task-related neuronal assemblies are more strongly reactivated in PFC during aSWR after learning

Learning-dependent reorganization of cortical circuits during memory consolidation has previously been linked to activity during hippocampal SWRs (Peyrache et al., 2009) and reactivation of spatial information in PFC during aSWRs has been reported by several groups (Kaefer et al., 2020; Maggi et al., 2018; Shin et al., 2019). aSWRs have additionally been implicated in the planning of goal-directed behavior (Ólafsdóttir et al., 2018). Therefore, we wondered if we could find evidence for reactivation of task-related neural assemblies during aSWRs occurring during inter-trial intervals of the conditioning task. To test this, we first detected the presence of neuronal cell assemblies in concatenated trial activity (Lopes-dos-Santos et al., 2013) and then checked the reactivation strength of these task related assemblies during aSWRs in CA1 and PFC. We found that reactivation of task-related assemblies in PFC increases significantly over the course of learning during hippocampal aSWRs (Figure 5A & 5B) (Wilcoxon rank sum test; p<0.05). This was true for assemblies defined during trials as well as during intertrial intervals (Figure S5.1). In CA1, on the other hand, reactivation strength remained constant from pre- to post-learning sessions (p=0.337). The frequency of aSWR occurrences remained constant between pre and post learning sessions (Pre n=24, 0.08 Hz; Post n=38, 0,09 Hz; Wilcoxon rank sum test; p=.2)(Figure S5.2A and S5.2B). aSWR rate decreased during trials and was at its lowest during reward consumption after CS+ trials (Figure S5.2C). Average aSWR rates independent of task stage slightly increased from the beginning to the end of each session (n=62; 0.06 Hz to 0.1 Hz, Wilcoxon rank sum test; p<0.01).

**Figure 5.**
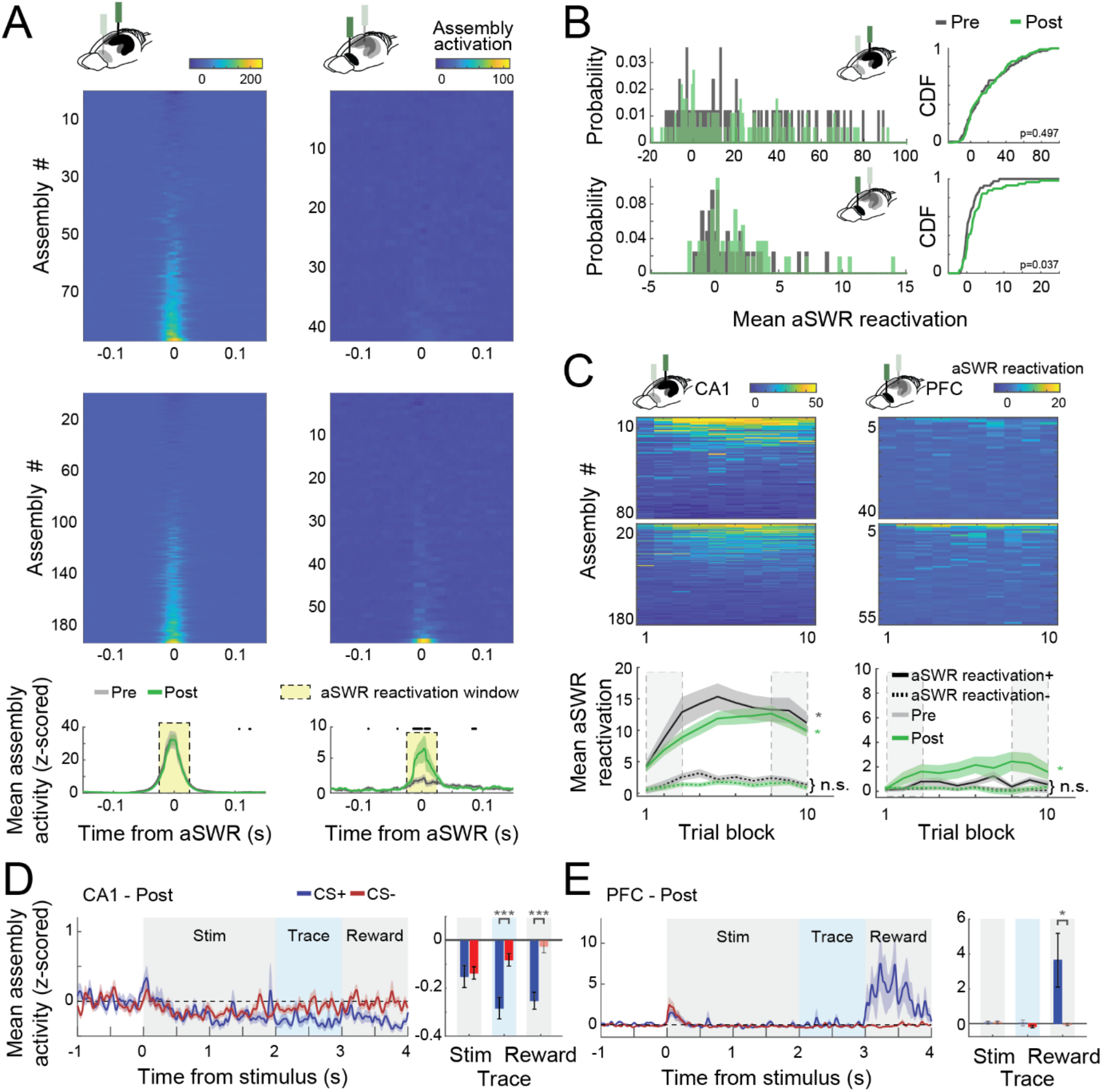
CA1 and PFC cell assemblies show different aSWR reactivation dynamics. **A)** Average (z-scored) assembly activation triggered by aSWR occurring in the inter-trial intervals for CA1 and PFC, Pre and Post learning sessions (Top). Mean aSWR-triggered activation over all the assemblies for Pre and Post sessions for each area. Shaded areas represent the SEM. Black dots represent windows in which Pre and Post assembly activity were statistically different (Wilcoxon rank sum test; p<0.05). Notice the higher aSWR triggered activation of assemblies in PFC in Post sessions. **B)** Histogram (left) and cumulative distribution function (CDF; right) of the mean assembly activity on the reactivation window denoted in A. P-values refer to a two-sample Kolmogorov-Smirnov test between Pre and Post distributions. **C)** Average aSWR reactivation of each assembly per session (Top). Sessions were divided into 10 blocks of equal trial length. Mean aSWR reactivation of all positively (reactivation+) and negatively (reactivation-) reactivated assemblies. Asterisks refer to Wilcoxon signed-rank test performed between the first and last three trial-blocks (dashed rectangles) of each area/learning condition (n.s.: non-significant; *p<0.05; **p<0.01; ***p<0.001) and shaded areas represent SEM. Note the evident increase in CA1 aSWR assembly reactivation across the session in both Pre and Post sessions for positively modulated assemblies (reactivation+). **D)** Mean (z-scored) assembly activity triggered by the stimulus onset for the 25% most strongly aSWR-reactivated assemblies in CA1 (Left). Average of the traces over each trial period is shown for CS+ and CS- (Right). Notice the initial decrease of assembly activity in CA1 during the stimulus and the posterior separation between CS+ and CS-. **E)** The same as in D, but for PFC assemblies. Note the difference between CS+ and CS-assembly activity during the reward period. Asterisks refer to a Wilcoxon signed-rank test comparing CS+ and CS- (*p<0.05; **p<0.01; ***p<0.001). Error bars refer to SEM and darker bars denote mean assembly activity significantly different from zero (p<0.05; t-test).

**Supplementary Figure 5.1.**
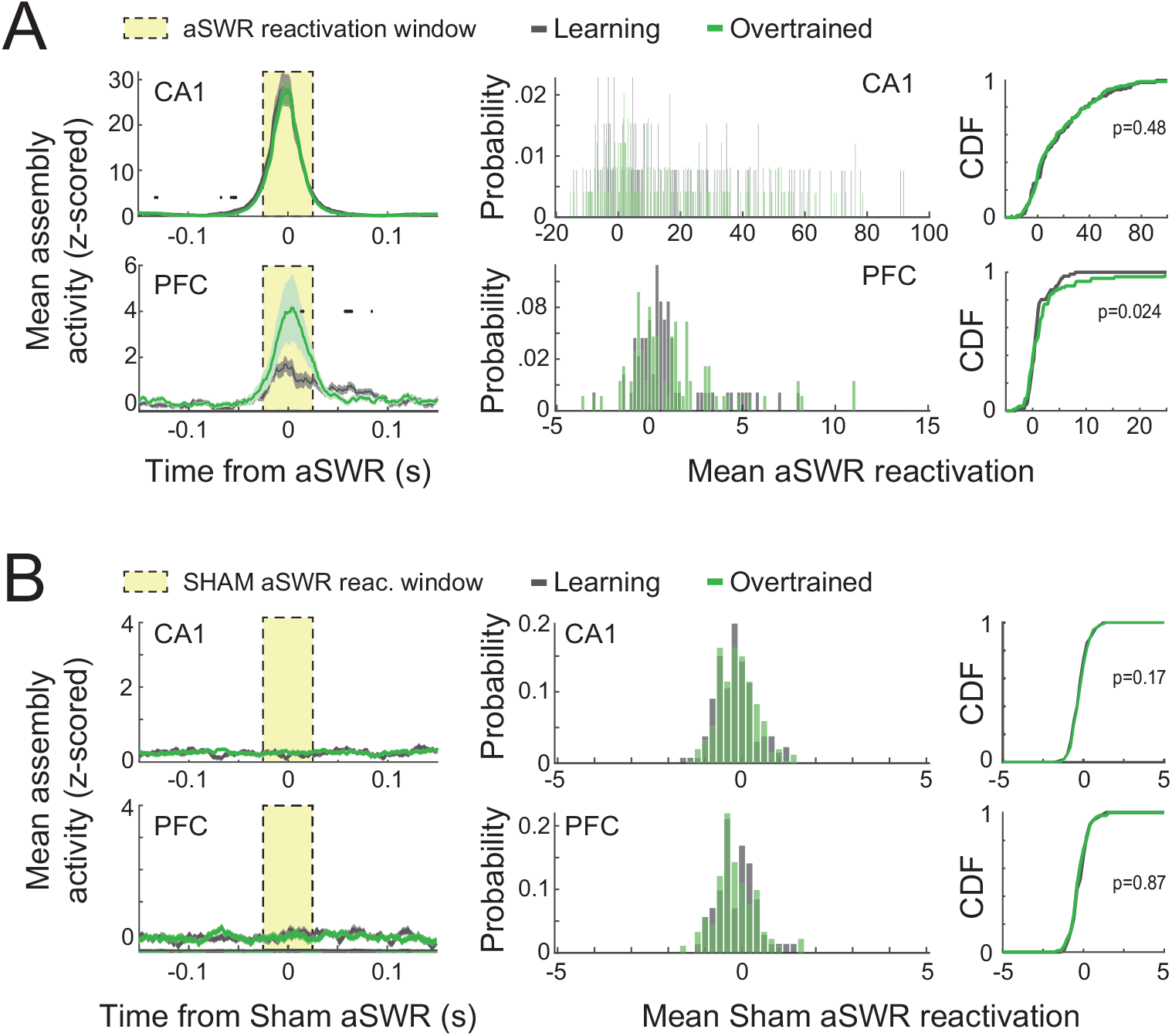
aSWR reactivation of assemblies detected during inter-trial intervals. **A)** (Left) Mean reactivation around aSWRs of assemblies detected during the inter-trial intervals (excluding aSWR events) for Pre and Post learning sessions. (Middle) Histogram of mean assembly aSWR-reactivation on the reactivation window (yellow rectangle) for Pre and Post learning sessions. (Right) Cumulative distribution of mean assembly aSWR-reactivation. P-values refer to a two-sample Kolmogorov-Smirnov test between Pre and Post distributions. **B)** Similar to A, but using sham aSWR times to compute the average reactivation. (aSWR events were randomly shifted by ∼200 ms).

**Supplementary Figure 5.2.**
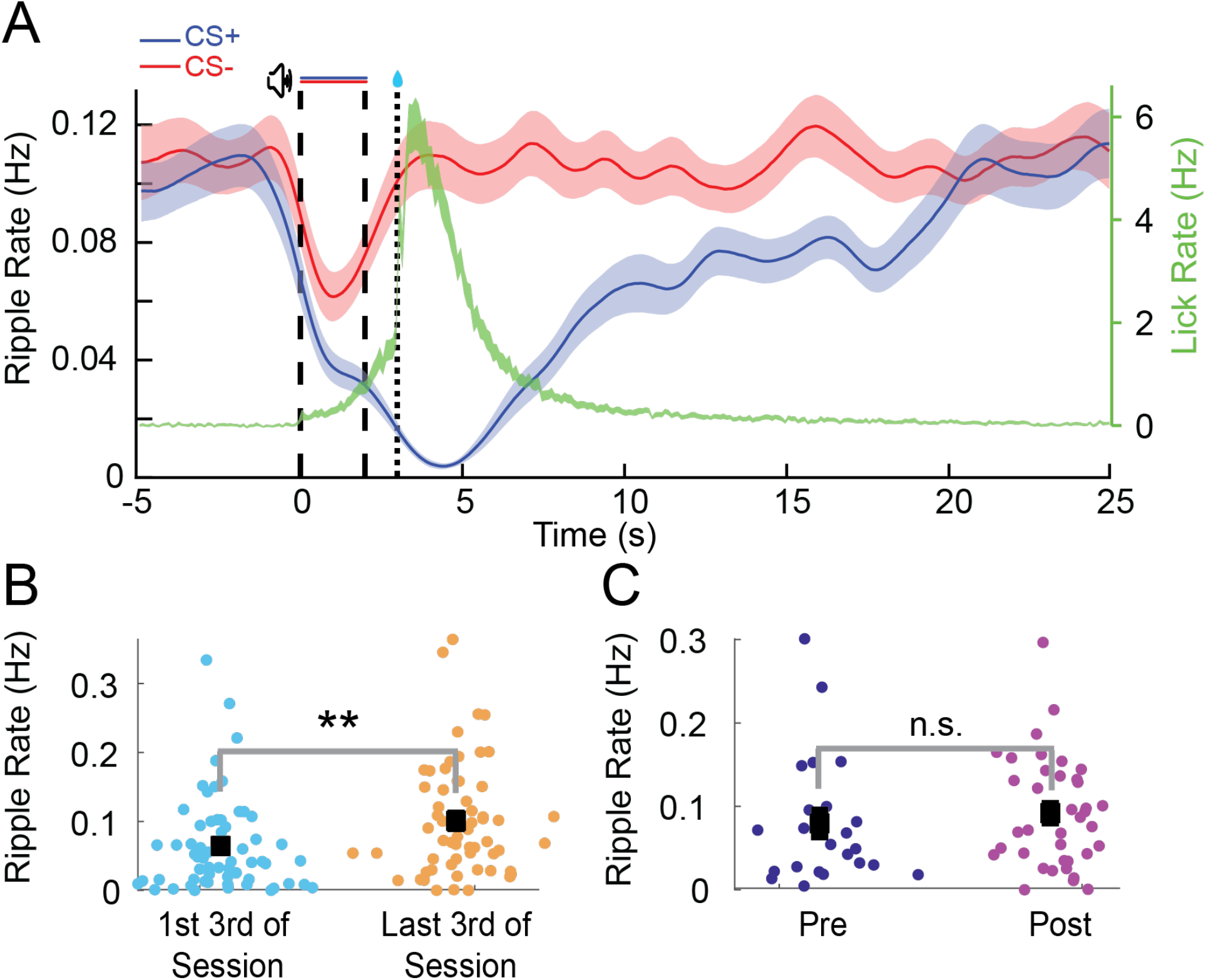
Distribution of awake Sharp Wave Ripples during trace-conditioning. **A)** Ripple rate during CS+(blue) and CS-(red) trials across all conditioning sessions. Average lick rate during CS+ trials is overlaid in green (Shaded areas indicate SEM). **B)** Average aSWR rate increases from early to late within individual sessions. **C)** Average aSWR rate does not change between pre and post learning sessions.

**Supplementary Figure 5.3.**
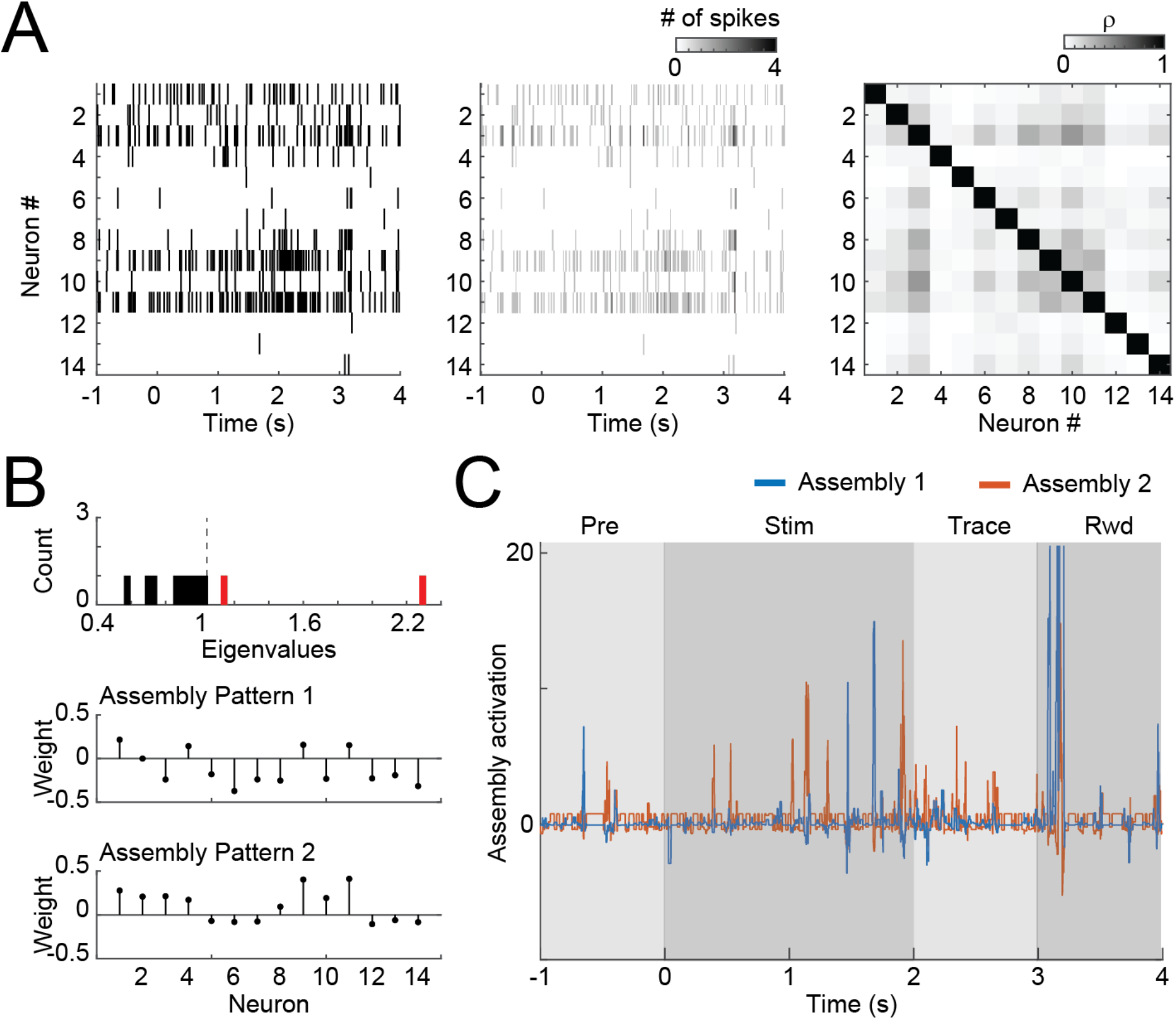
Detecting cell assemblies in neural populations. **A)** The rastergram (left) of each trial was computed and binned in 20-ms-bins with no overlap (middle). After concatenating the activity of all trials, the activity of each neuron was z-scored and the correlation matrix was computed (right). **B)** The eigenvalues of the correlation matrix were then computed and compared to the analytical (Marchenko-Pastur) distribution to estimate the amount of assembly patterns present in the data (top). After that, independent component analysis was used to extract the assembly patterns (bottom). **C)** The patterns in B were then used to project the assembly activity during the trial, using 20 ms bins with steps of 1 ms.

### Task-related assembly reactivation strength increases during individual sessions in CA1

We next analyzed how reactivation strength changed over time within individual sessions. We found that in CA1 assembly reactivation strength per aSWR increases gradually over the course of individual pre-learning sessions for positively modulated assemblies (reactivation+) as well as for negatively modulated assemblies (reactivation-) during aSWRs (Figure 5C). This effect was also present in post-learning sessions for the reactivation+ assemblies (One-way ANOVA; CA1 pre+: p<0.01, pre-: p<0.05, pos+: p<0.001, post-: n.s.). In PFC assembly reactivation strength slightly increased for positively modulated assemblies in post learning sessions (Figure 5C).

We then sought to determine what type of task specific information the most strongly SWR-reactivated assemblies represent during the conditioning task in post learning sessions. In CA1, we found that the 25% most reactivated assemblies are suppressed during the trace and reward period after CS+ trials compared to CS-trials (Figure 5D) (Wilcoxon signed-rank; p<0.001). In PFC, on the other hand, the 25% most reactivated assemblies responded strongly during the reward period (Figure 5E) (t-test, compared to pre-trial baseline; p<0.05). Those effects were not observed for the 25% least reactivated cells.

### CS+ responsive assemblies are preferentially replayed in CA1 during aSWRs

Given that assemblies detected during the task and the intertrial intervals became both reactivated during aSWR, we next asked whether we could find additional evidence for a prioritized reactivation of assemblies that carry specific information about CS sounds during the trials. To this end, we first computed the average activity of each assembly for CS+ or CS-stimuli and computed a trial-type modulation score, defined as the average assembly activation during CS+ trials subtracted by the average activation during CS-trials (stimulus and trace periods; Figure 6A). Then, for each session, we selected the most positively modulated (i.e., CS+ activity higher than CS-), the most negatively modulated (i.e., CS-activity higher than CS+), and the least modulated, control assembly. Because we found that most assemblies in CA1 are suppressed during the stimulus (as are the firing rates), we term assemblies by whether they were more suppressed during CS+ or CS-stimuli. We found that, for CA1, CS+ suppressed assemblies in both stimulus and trace periods (i.e., assemblies that are more suppressed during CS+ trials than CS-trials) were more reactivated during aSWRs (Figure 6B) compared to CS-suppressed assemblies and control, non-modulated assemblies. This effect was confirmed by the presence of a negative correlation between the trial-type modulation score and the average aSWR reactivation of each assembly (Figure 6C).

**Figure 6.**
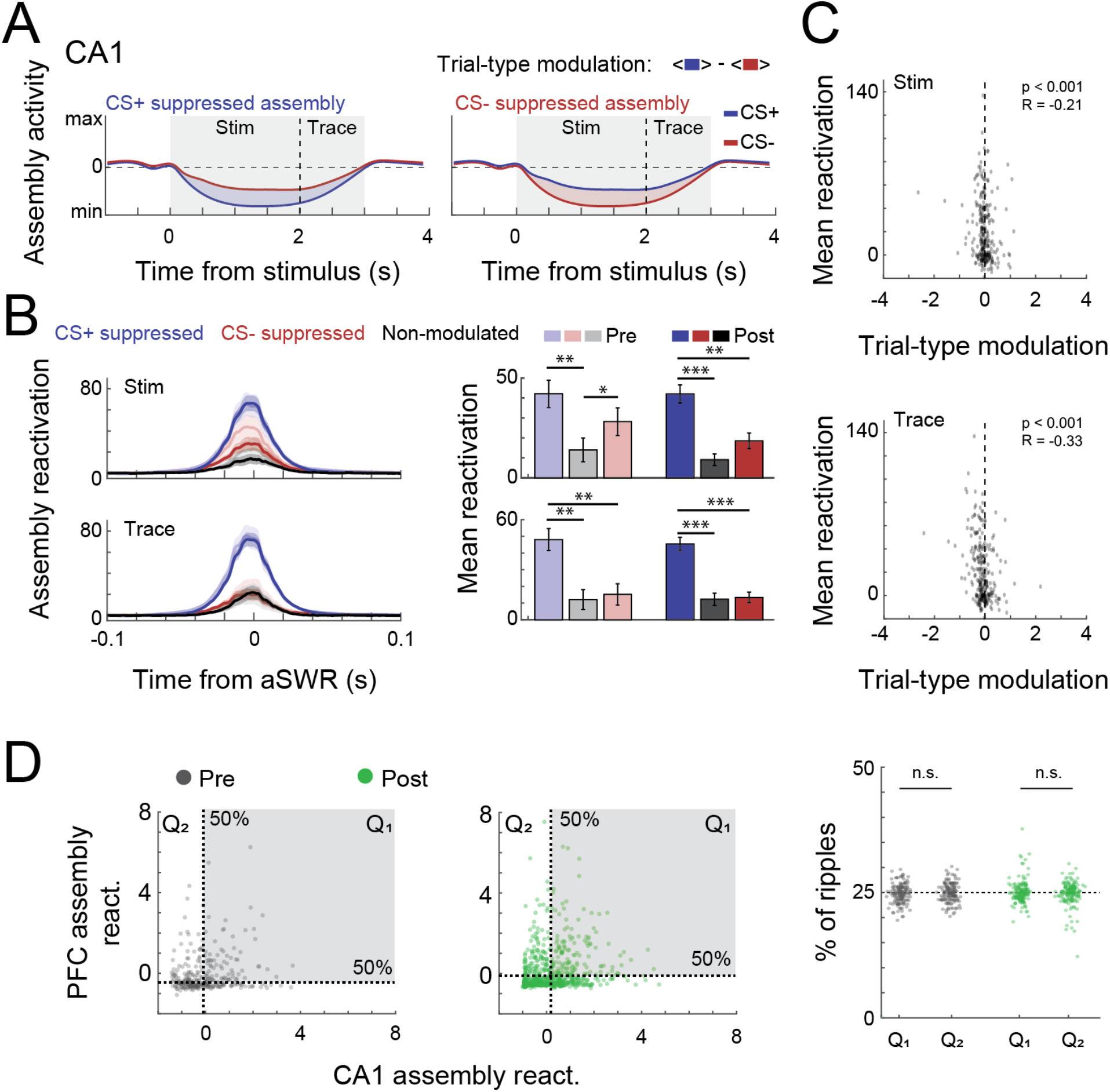
Trial-type modulation and PFC coactivation of CA1 assemblies. **A)** Schematic representation of trial-type modulation scores, CS+ and CS-suppressed assemblies. The modulation score was defined as the difference between average assembly activation on CS+ and CS-trials during a specific period. **B)** Mean aSWR reactivation of CS+ suppressed, CS-suppressed and non-modulated assemblies pre and post learning over time (left) and within 50 ms window around ripples (right). Error bars denote SEM (*p<0.05; **p<0.01; ***p<0.001). Scatter plot and Pearson’s correlation values between trial-type modulation score and average aSWR reactivation for all CA1 assemblies (pre and post learning). Notice the stronger reactivation of negatively modulated assemblies (CS+ suppressed). **D)** (Left) Example of joint reactivation for two pairs of CA1-PFC assemblies. Quadrants were defined using the median aSWRs reactivation of each area and the proportion of reactivations in each quadrant was computed. (Right) Percentage of ripple reactivations in 1st and 2nd quadrants defined in left for all possible combination of assembly pairs (Wilcoxon signed-rank test).

CS-suppressed assemblies were also more strongly reactivated than control assemblies in pre-learning sessions (p<0.05, Figure 6B) and CS-assemblies that were more suppressed during stimulus compared to trace period also reactivated more during aSWR (Supp Figure 6 C). Together this indicates that CS-coding assemblies are also differentially modulated during aSWR.

In PFC we did not observe preferential reactivation of CS+ or CS-specific assemblies (Supp. Figure 6).

**Supplementary Figure 6.**
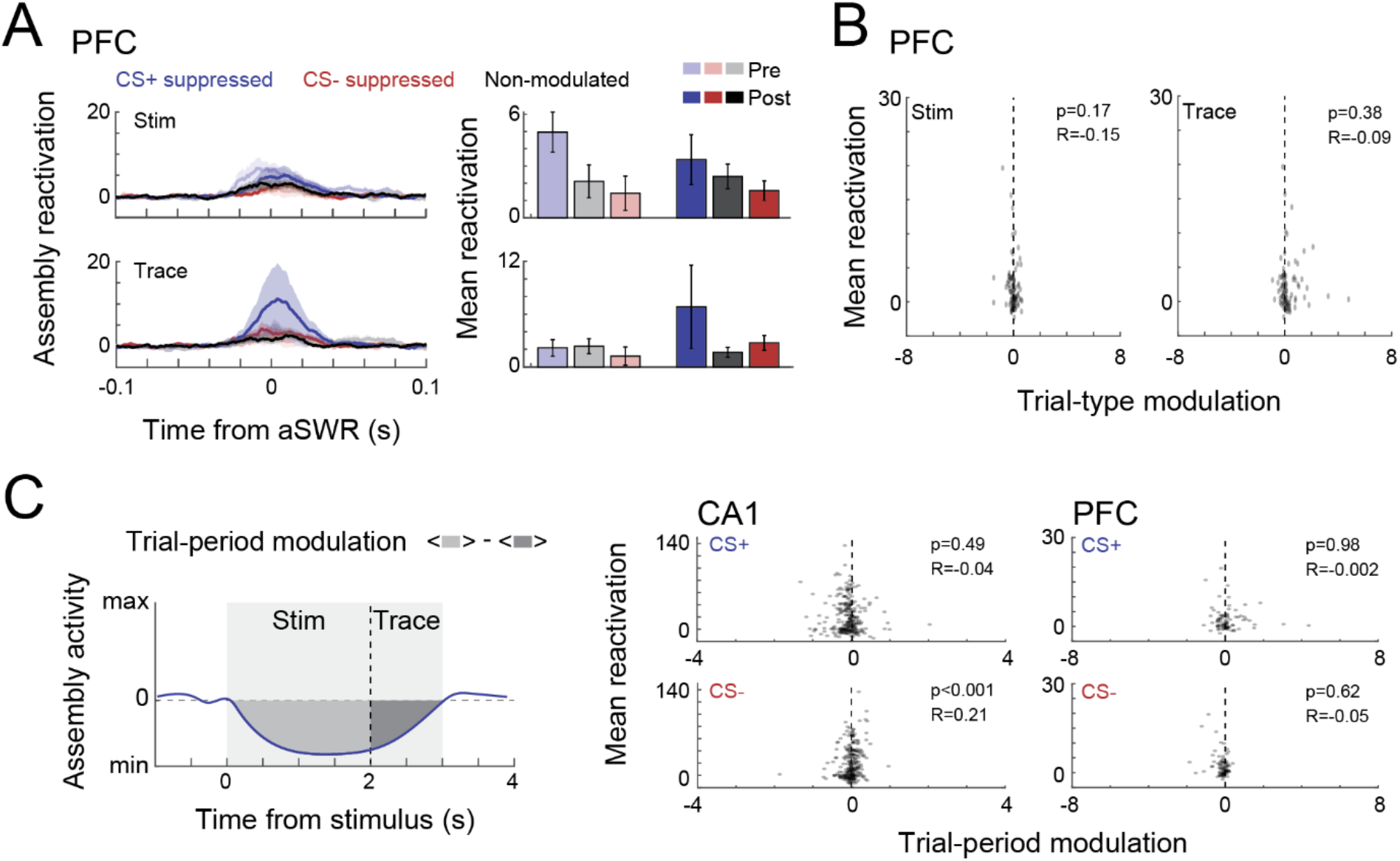
Trial-type modulation of PFC assemblies and trial-period modulation. **A)** Mean aSWR reactivation of CS+ suppressed, CS-suppressed and non-modulated assemblies in PFC over time (left) and within 50 ms window around ripples (right). Error bars denote SEM. **B)** Scatter plot and Pearson’s correlation values between trial-type modulation score and average aSWR reactivation for all CA1 assemblies (pre and post learning). **C)** (Left) Schematic representation of trial-period modulation scores. The trial-period modulation score was defined as the difference between average assembly activation on stimulus and trace periods in CS-trials. (Right) Scatter plot and Pearson’s correlation values between trial-period modulation scores and aSWRs reactivation for assemblies in CA1 and PFC.

Finally, we wondered whether CA1 and PFC assembly reactivation is coordinated during aSWRs. Coordinated reactivation of task relevant information during aSWR has previously been found in the CA1-PFC circuit during some spatial navigation tasks but not during others(Kaefer et al., 2020; Shin et al., 2019). To check whether CA1 and PFC assembly reactivation is coordinated during aSWR during trace conditioning, we computed how often high aSWR reactivation events co-occurred in the two areas (Figure 6D). In line with previous reports (Kaefer et al., 2020), we did not find evidence for coordinated reactivation of CA1 and PFC assemblies during aSWRs.

## Discussion

This study characterizes changes in neural activity in the CA1-PFC network while mice learn to use predictive sounds to anticipate future rewards. We show that activity in both areas is strongly shaped by learning and that task specific information is reactivated in a complex pattern across CA1 and PFC during aSWR.

While CA1 and PFC are highly active during aversive eyeblink trace conditioning, evidence for a similar involvement during appetitive trace-conditioning had been missing. In fact, several previous studies have pointed out differences in the mechanisms underlying both types of learning (Pezze et al., 2017; Thibaudeau et al., 2007).

Despite these differences, we found that single cells in CA1 and mPFC during appetitive trace-conditioning behave similarly to what had previously been reported during aversive trace-conditioning. Both areas display long lasting sustained activity that bridges the temporal gap between CS+ offset and reward delivery. In both areas, these sustained responses are composed of a mix of Trace-Up and Trace-down cells, i.e., cells that display sustained excitation and inhibition respectively. In CA1, higher numbers and stronger inhibition of Trace-Down cells result in overall suppression of the entire area during the trace period, while in PFC higher numbers and stronger activation of Trace-Up cells resulted in overall excitation.

Similar to our study, abundant Trace-Down like responses and spares Trace-Up like responses have been described during aversive trace-conditioning in CA1 (Hattori et al., 2015; McEchron and Disterhoft, 1997). This distinct pattern of mostly inhibition mixed with sparse excitation has been hypothesized to increase the signal to noise ratio to more efficiently propagate the signal of Trace-Up cells to downstream areas (Hattori et al., 2015) Yet, it is also conceivable that Trace-down cells participate in an independent form of coding. Inhibition in CA1 might for example play an active role in suppressing well expected incoming stimuli, i.e. reward delivery (Bastos et al., 2012; Rummell et al., 2016; Stachenfeld et al., 2017).

In PFC, responses during appetitive trace-conditioning are also similar to what has previously been found during aversive trace-conditioning. Specifically, higher numbers and stronger excitation of Trace-Up cells have also been found in rat PFC and parts of rabbit PFC during aversive trace-conditioning (Hattori et al., 2014; Takehara-Nishiuchi and McNaughton, 2008). A learning dependent reduction in responses to CS-like pseudo conditioning stimuli have also previously been described in PFC (Hattori et al., 2014; Takehara-Nishiuchi and McNaughton, 2008; Weiss and Disterhoft, 2011).

In combination, this suggests that sparse excitation with strong surrounding inhibition in CA1 and mostly excitation in PFC are two general coding principles employed to bridge the temporal gap between a salient cue and a behaviorally relevant event, independently of the appetitive or aversive nature of the event and the specific anticipatory action that it requires. We furthermore observed high frequency CA1-PFC coherence was specifically increased during CS+ trials. Increased synchronization between both areas during spatial working memory tasks have previously been reported predominantly in the Theta frequency (4-12 Hz) (Battaglia et al., 2011; Benchenane et al., 2010; Jones and Wilson, 2005). However, during trace-conditioning mice were mostly stationary and did not display strong hippocampal theta oscillations and high frequency coherence therefore likely results from a different underlying mechanism.

Despite both areas being highly engaged in the task and encoding trial specific information on a trial-by-trial level, we did not find any evidence for task-specific communication on the single cell level. CA1 and PFC therefore either process conditioning trials in parallel rather than in series or rely on intermediate structures (e.g. entorhinal cortex) for effective communication (Insel and Takehara-Nishiuchi, 2013).

Lastly, we found that cell assemblies in CA1 and PFC that are responsive during classical conditioning also strongly reactivate during awake hippocampal Sharp-Wave Ripples (aSWRs). Earlier reports have found coordinated activity in CA1 and PFC to be elevated during aSWRs and to change over the course of learning of a spatial memory task (Kaefer et al., 2020; Shin et al., 2019; Shin and Jadhav, 2016; Tang et al., 2017) However, physiological evidence for reactivation of neural assembly patterns during aSWRs in a non-spatial task had been missing. Our data now provides this missing link and adds to the notion that reactivation of task-related patterns of neural activity during aSWRs plays a general role during learning and action planning (Joo and Frank, 2018; Ólafsdóttir et al., 2018). Notably, we observed a fast increase in reactivation strength within CA1 as individual training sessions progressed and a slow increase in reactivation strength in PFC from pre-learning to post-learning sessions. This is well in line with the idea that both areas support learning on different timescales, with the hippocampus adapting fast to new experiences and the prefrontal cortex adapting more slowly to behaviorally important variables that remain stable over time (McClelland et al., 1995; Takehara-Nishiuchi and McNaughton, 2008; Takehara et al., 2003). Assembly reactivation in CA1 was furthermore task specific during trace-conditioning Assemblies that were suppressed during and after CS+ sounds became most strongly reactivated. CS-suppressed assemblies were also but less strongly modulated during aSWRs in CA1.

This preferential reactivation of CS suppressed assemblies during aSWR, provides additional evidence that suppression in CA1 during CS+ trials plays a pivotal role during trace conditioning and might be relevant to actively encode stimulus identity or to predict upcoming task events.

In contrast, aSWRs-reactivated assemblies in PFC strongly encode reward and trial unspecific information which could reflect a reactivation of learned, context dependent rules that has previously been described during slow wave sleep SWRs (Peyrache et al., 2009). We did not observe that assemblies in CA1 and PFC co-reactivated during aSWR. This is well in line with previous reports (Kaefer et al., 2020) and suggest that both areas can process some forms of task related information independently during aSWRs.

## Material and Methods

### Animals

For this experiment, we used a total of 17 male C57/Bl6j mice. The animals were obtained at 10-13 months of age from Charles River Laboratory and all experiments were performed within two months after delivery. 7 animals were used for silicon probe recordings from dorsal hippocampus area CA1, 6 animals were used for recordings from PFC and an additional 4 animals were used for combined silicon probe recordings from CA1 and PFC during the AATC task. All animals were group housed until the first surgery after which they were individually housed to prevent damage to the implants. Throughout the experiment the animals were maintained on a reversed 12-hour light/dark cycle and received food and water ad libitum until we introduced food restriction two days after the first surgery. All experiments were performed during the dark period. This study was approved by the Central Commissie Dierproeven (CCD) and conducted in accordance with the Experiments on Animals Act and the European Directive 2010/63/EU on animal research.

### Surgical preparation for head-fixed electrophysiological recordings

Animals were anesthetized using isoflurane (1–2%) and placed in a stereotaxic frame. At the onset of anesthesia, all mice received subcutaneous injections of carprofen (5 mg/kg) as well as a subcutaneous lidocaine injection through the scalp. The animals’ temperature was maintained for the duration of the surgical procedure using a heating blanket. Anesthesia levels were monitored throughout the surgery and the concentration of isoflurane was adjusted so that a breathing rate was kept constant at around 1.8 Hz. We exposed the skull and inserted a skull screw over the cerebellum to serve as combined reference and ground for electrophysiological recordings. We then placed a custom made, circular head-plate for head-fixation evenly on the skull and fixated it with dental cement (Super-Bond C&B). For CA1 recordings, a craniotomy was performed over the left hippocampus -2.3 mm posterior and +1.5 mm lateral to Bregma and for PFC recordings a craniotomy was performed over left frontal cortex at +1.78mm anterior and +.4mm lateral to Bregma. The exposed skull was covered with a silicon elastomer (Body Double Fast, Smooth-on) until the first recording. All mice were given at least 2 days to recover from the surgery.

### Head-fixed virtual reality setup

The head-fixed virtual reality setup consisted of two rods that were screwed onto either side of the implanted head-plate and fixated the mice on top of an air-supported spherical treadmill. The motion of the treadmill was recorded using an optical mouse and transformed into movement along a virtual linear track designed with the Blender rendering software. The virtual track was then projected through a mirror into a spherical screen surrounding the head-fixed animal on the treadmill (Schmidt-Hieber and Häusser, 2013)

(https://github.com/neurodroid/gnoom). While in head-fixation the animals received soy milk as reward which was delivered through a plastic spout that was positioned .5cm anterior to the lower lip. Licks were detected with an infrared beam-break sensor that was positioned right in front of the spout.

All animals were slowly habituated to head-fixation by placing them in the setup for at least 2 days of 3×10 minute sessions during which they received about 50 rewards, totaling to about .2ml of soy milk. During the habituation, we started to food restrict the animals to around 90% of initial body weight to motivate better task performance.

In all cases, the food restricted animals started to lick off the soymilk reward reliably within the first 6 habituation sessions.

### Behavioral training

The AATC task required the animals to associate a 2-second-long CS+ sound with a droplet of soymilk reward (∼5 microliter), delivered after 1 second of silence, the so-called “trace period” while ignoring a CS-control sound. We interleaved the CS presentations randomly every 30-45 seconds. For the two sounds, we choose a 3000 kHz continues pure tone and a 7000 kHz tone pulsating at 10hz. We counter-balanced the CS+ and CS-sounds evenly between animals throughout the experiment.

As the main behavioral outcome measure, we detected the licks of the animals with an infrared beam-break sensor that was mounted in front of the reward spout. Each training session ended as soon as the animals received 50 rewards. We repeated the experiment for at least 10 days.

During these behavioral training sessions, the head-fixed animals could freely run on the linear track in the virtual reality which was otherwise not correlated with the AATC task.

We performed acute silicon probe recordings from the dorsal CA1 area and/or PFC of head-fixed mice during the first 2 days of the AATC task as well as from day 6 onwards for as long as we were able to achieve stable recordings. We then classified recording sessions from the first 2 days of training as pre-learning (Pre) and recording sessions from day 6 onwards and with a significant increase in anticipatory licks as post-learning (Post).

### Acute electrophysiological recordings during AATC tasks

At the start of each recording session, we placed the mice in head-fixation and removed the silicon elastomer cover to expose the skull. We then used a micromanipulator (Thorlabs) to acutely insert a 128-channel silicon probe into the middle of the previously prepared craniotomy above PFC and/or CA1. For PFC recordings, we then slowly lowered the recording electrode to -2.0mm ventral to Bregma. For CA1 recordings, we continuously monitored the local field potential (1-600Hz), ripple frequency signal (150-300Hz) and spiking activity (600-2000Hz) during the insertion process and tried to positioned our electrode in a way that the strongest ripple amplitude and spiking activity was 200-250µm from the base of the probe and 470-520µm from the tip. In this way, we were able to cover most of dorso-ventral extend of CA1 with our recording electrode.

Electrophysiological signals were filtered between 1 and 6000 Hz, digitized at 30 kHz using 2 64 channel digitizing heads-stages (RHD2164 Amplifier Board, Intan Technologies) and acquired with an open-ephys recording system. After each recording session, we retracted the silicon probe and placed a new silicon cover on the skull before releasing the animals back to their respective home cages.

### Behavioral Data Analysis

For every training session and each animal, we compared the change in lick-rate between CS+ vs CS-trials. In short, for each trial we took the sum of all licks during the trace period and subtracted the sum of all licks during the baseline period (−1s to onset of CS). We then computed a T-test between the change in lick rate for all CS+ vs all CS-trials and defined the animal to have learned (post-learning sessions) if this comparison showed a significant difference in lick rate between the two conditions.

### Neural Data Analysis

To identify single unit activity, the raw voltage signal was automatically spike sorted with Kilosort (Pachitariu et al., 2016) (https://github.com/cortex-lab/Kilosort) and then manually inspected and curated with the ‘phy’ gui (https://github.com/kwikteam/phy). All following analysis was performed using custom written Matlab scripts.

### Single Cell responses

For each unit, we binned the single cell spiking data (25ms), smoothed the data with Gaussian-weighted moving average filter (25 Bins) and computed the Peri-Stimulus-Time-Histograms (PSTH) for CS+ and CS-trails.

To assess evoked responses to CS+ and CS-we calculated the Z-scored firing rates in the first 350ms (post-learning CS) post stimulus interval for each cell and compared population responses in this time window.

To assess lick related activity, we first defined lick onset as the first lick after CS+ sound onset but before reward delivery that was followed by at least 3 licks within the next second. We then defined single cells to be lick-responsive if the average firing rate during the time window around the 1st lick (−250ms to +250ms) was increased by at least 1 standard deviation or decreased by at least 1 standard deviation from pre-trial baseline (−1000 - CS+ onset) To assess trace period activity, we defined single cells to be trace responsive if the average Z-scored firing rate during the 1 second trace period increased by at least 1 standard deviation or decreased by at least 1 standard deviation from baseline during either CS+ or CS-trials. To quantify group differences, we performed a Kruskal-Valis (pre vs post learning) using average S-scored responses during the trace period.

For analysis of learning dependent changes in reward evoked activity, we calculated the Z-scoredfiring rates during the reward response window (reward delivery-reward delivery+500ms).

### Population rate vector analysis

To analyze the CA1 and PFC population response during the trace period, we first computed the average CS+ and CS-PSTHs for all simultaneously recorded non-lick cells in 25ms bins for every session and smoothed the data with Gaussian-weighted moving average filter (25 bins). For visualization purposes, we then computed the first 3 principal components of the resulting matrices CS+ and CS-PSTHs and plotted the resulting 3-dimensional vectors. To quantify the difference in population activity between CS+ and CS-trials, we calculated the Euclidean distance between CS+ and CS-in n-dimensional space (n=number of simultaneously recorded non-lick cells) for each bin and averaged across sessions.

In order to predict trial identity by trace period population firing rates using a support vector machine classifier, we first calculated the trial-by-trial firing rates for all simultaneously recorded cells during the trace period. We then split all trials of a single recording session into 20 equal partitions and used 19 of these partitions to train the support vector machine classifier (fitcsvm function, MATLAB). We then tested the classifier performance on the 20^th^ partition and repeated this process for all other partitions (20fold cross validation). Classifier performance above chance was determined by comparing the average prediction accuracy across all sessions against chance level using a Wilcoxon sign rank test.

### Reduced Rank Regression of CA1-PFC single cell interactions

To investigate CA1-PFC interaction on the single cell population level, we used reduced rank regression (RRR) to assess how well the activity of a sampled population in one of the areas (source area) can be explain another (disjoint) sampled population in the same or in a separate target area, through a simplified, low-dimensional linear model. (Semedo et al., 2019). Briefly, for a given session, we first subsample (without replacement) the population of each region in two equally-sized sets of source and target neurons, so that all four sets had the same number of neurons. We then used a 10-fold cross validation scheme to compute the RRR (i.e., fitting the target population activity using the source population) using multiple rank values. The performance of each model was computed using the relative amount of variance explained by the model (R2). We then selected the first model which had mean performance within 1 SEM of the best model, using its rank as the number of predictive dimensions (Figure 7B). This procedure was repeated for 10 different subsamples and the performance and number of predictive dimensions of each session was computed via averaging across sub-samples. We also compared the performance of the full regression model (in which all the ranks were used) to control for different dimensionality of the RRRs in the two areas. In this particular case, we added L1 regularization, and chose the best model (highest average performance over cross-validation) among different ridge parameter values and measured the MSE between estimated and real activity (Supp. Figure 7A).

### Coherence analysis

To assess coherence between mPFC and CA1 during the AATC task, we first selected the CA1 recording channel with the strongest aSWR amplitude (see below) as well at the central channel of the mPFC recording electrodes and down-sampled the raw voltage singles to 2000Hz. Coherence was then analyzed with multi-taper Fourier Analysis (<u>Mitra and Pesaran, 1999</u>), using the Chronux MATLAB toolbox (http://www.chronux.org).

### Reactivation during awake Sharp-Wave Ripples

In order to detected awake Sharp-Wave Ripples during the inter-trial periods of the AATC task, we first filtered the local field potential at the top half of our recording electrode (16 channels in total, 1 channel at each depth, 320µm spread around CA1 cell layer) between 150-300Hz and used common average reference to exclude artefacts affecting all channels. We then identified the recording depth with the strongest average ripple power and used the ripple band signal on this channel for further analysis. For each recording session, we visually inspected the ripple band signal and manually set a low-cut threshold for ripple detection (100 µV in most cases) and a high-cut threshold for artefact rejection (500µV in most cases). We furthermore, excluded threshold crossings within 200ms of each other as well as within 200ms of any licking activity.

To compute task related cell assemblies in CA1 and PFC, we first binned the spikes of each trial (−1 to 4 s from stimulus onset) into 20 ms bins. Then, we used independent component analysis (ICA) to find the co-activation patterns as described previously (Lopes-dos-Santos et al., 2013). The number of assemblies was defined by the eigenvalues of the cross-correlation matrix that were above the analytical Marcenko Pastur distribution (Lopes-dos-Santos et al., 2013). We then projected the neural activity onto each of the assembly patterns (using the same 20 ms bins, with overlap) and computed the mean assembly activity triggered by the stimulus, animal licks and hippocampal SWR, normalizing it with the z-score transformation. Normalization was done using the z-score transformation, which in the case of stimulus-triggered assembly activity only used pre-stimulus period (of both CS+ and CS-) as baseline (−1000 - 0 s) and in case of ripples-triggered activity only used the period outside the ripple center (50 ms window) as baseline. At last, we defined the assembly reactivation window on SWR as the average z-scored assembly activity in the 50 ms window centered on the aSWR. We then divided each session in 10 equally long trial blocks, and investigated the reactivation strength of positively (reactivation+) and negatively (reactivation-) modulated assemblies over the course of the session. Statistical comparisons between pre and post (normalized) mean assembly activity was done using a Wilcoxon rank sum test, while comparisons of reactivation within a session was done comparing the reactivation in the first three and last blocks using a Wilcoxon signed-rank test. In Figure 5D-E traces were smoothed using a 100 ms Gaussian window. Comparisons between CS+ and CS-stimuli were done using a Wilcoxon signed-rank test, while comparisons with baseline (pre-stimulus period) were done using a t-test.

### Trial-type encoding assembly reactivation during aSWR

For each assembly a trial-type modulation score (Figure 6A) was computed defined as the average assembly activation (in a given period) during CS+ trials minus the average activation during CS-trials. Similarly, a trial-period modulation score was defined as the average assembly activation during stimulus minus the average assembly activation during trace (Supp. Figure 6A). For trial-period modulation scores, only CS-trials were used. In Figure 6B and Supp. Figure 6B only the assemblies with the lowest (CS+ suppressed), highest (CS-suppressed) and least (non-modulated) score values of each session were chosen.

### CA1-PFC assembly co-activation during aSWR

To assess the coordination of CA1 and PFC assembly reactivation (during aSWRs), we counted how often each simultaneously recorded assembly pair (1 CA1 and 1 PFC assembly) reactivated together above the median in both areas and then compared the percentage of coincident high reactivations (Q1) with the percentage of high and low reactivations in CA1 and PFC respectively (Q2).

### Histology

At the end of each experiment, mice were perfused with 4%PFA and brain sections (100µm) were examined with light microscopy to confirm electrode placement in CA1 and mPFC.

## Acknowledgments

Funding was provided by a German Studiensstiftung fellowship (to JK), by the Dutch NWA “Bio-Art” project (to FPB), and by the NWO Top-grant no. 612.001.853 (to FPB)

## Competing interests

No competing interests declared.

## Notes

### Competing Interest Statement

The authors have declared no competing interest.

